# Comparative genomics of apomictic root-knot nematodes: hybridization, ploidy, and dynamic genome change

**DOI:** 10.1101/136085

**Authors:** Amir Szitenberg, Laura Salazar-Jaramillo, Vivian C. Blok, Dominik R. Laetsch, Soumi Joseph, Valerie M. Williamson, Mark L. Blaxter, David H. Lunt

**Affiliations:** Evolutionary Biology Group, School of Environmental Sciences, University of Hull, Kingston upon Hull, UK; The Dead Sea and Arava Science Center, Microbial Metagenomics Division, Masada, Israel; Institute of Evolutionary Biology, School of Biological Sciences, University of Edinburgh, Edinburgh, UK; The James Hutton Institute, Invergowrie, Dundee, UK; Department of Entomology and Nematology, University of Florida, Gainesville, USA; Department of Plant Pathology, University of California, Davis, USA

**Author notes:** Correspondence: Amir Szitenberg.

**Keywords:** *Meloidogyne*, Genome evolution, Phylogenomics, Coverage ratio, Recombination

## Abstract

The Root-Knot Nematodes (RKN; genus Meloidogyne) are important plant parasites causing substantial agricultural losses. The Meloidogyne incognita group (MIG) of species, most of which are obligatory apomicts (mitotic parthenogens), are extremely polyphagous and important problems for global agriculture. While understanding the genomic basis for their variable success on different crops could benefit future agriculture, analyses of their genomes pose challenges due to complex evolutionary histories that may incorporate hybridization, ploidy changes, and chromosomal fragmentation. Here we sequence 19 genomes, representing five species of key RKN collected from different geographic origins. We show that a hybrid origin that predated speciation within the MIG has resulted in each species possessing two divergent genomic copies. Additionally, the MIG apomicts are hypotriploids, with a proportion of one genome present in a second copy, and this proportion varies among species. The evolutionary history of the MIG genomes is revealed to be very dynamic, with non-crossover recombination both homogenising the genomic copies, and acting as a mechanism for generating divergence between species. Interestingly, the automictic MIG species *M. floridensis* differs from the apomict species in that it has become homozygous throughout much of its genome.

## Introduction

The Root-Knot Nematodes (RKN; genus *Meloidogyne*) are among the world’s most destructive crop pests, causing very significant reduction in yields in nearly all major agricultural crops [1]. The most-studied species in this genus can be divided into three well-supported clades, and the tropical RKN species in Clade 1, especially the closely related *Meloidogyne incognita* group (MIG) species, are globally significant pests. They are found in agricultural areas on all continents that have mild winter temperatures [2], and have been highlighted as one of the most serious threats to temperate agricultural regions as climate change progresses [3].

Based on cytological examination of gamete development, most MIG nematodes have been determined to be mitotic parthenogens (apomicts), which do not undergo meiosis and reproduce asexually [4–7]. Although asexual organisms are often characterised as being less able to adapt to variable environments and interspecific competition than those with meiosis, the MIG apomicts are very successful, globally-distributed, highly-polyphagous crop pests [2]. Considerable variation in ability to break crop resistance and to reproduce on different crop species is observed both between and within species [2,8]. Despite their global importance relatively little information is available on genetic variation between and within species, or genomic diversity across pathogenicity groups and mating systems. Draft genomes are available for two MIG species: *Meloidogyne incognita* [9] and *Meloidogyne floridensis* [10]. A high quality genome assembly is available for the distantly-related *Meloidogyne hapla*, a facultative meiotic parthenogen from *Meloidogyne* Clade 2 [11]. It has been suggested on the basis of the two draft genome sequences that the MIG arose from a complex series of hybridisation events, and this hybrid origin is evident in the genome structure [10]. Genome-scale data are important if we are to accurately understand the complex origins and evolution of the MIG and their different abilities to increase in virulence on crops.

The MIG species have classically been differentiated using subtle morphological characters, isozymes, and host range [12–16]. The phylogenetic relationships between the closely-related MIG species have been difficult to unequivocally determine [4,17–20]. Nuclear genome sequencing has revealed that MIG species contain two very divergent copies of many loci, likely due to a past hybridisation event [21]. The different evolutionary histories of these copies, likely to have been brought together by hybridisation (*i.e.* they are homoeologs) compromise phylogenetic analyses and species identifications that use nuclear genome sequences. While mitochondrial DNA approaches can successfully discriminate some MIG species [20,22–24], there is little phylogenetic signal and these maternal-lineage mtDNA studies cannot report on hybrid origins. Large-scale sequencing of nuclear genomes from multiple species and isolates has the potential to provide a wealth of comparative data. Modern population genomic analysis techniques will then be available to discriminate closely related species, examine the agricultural spread and divergence of populations, more fully represent the MIG’s evolutionary history and phylogenetic relationships.

We have sequenced the genomes of 19 new *Meloidogyne* isolates from five nominal species, including 1 to 8 unique isolates of each species from diverse geographical locations. The new genomes include isolates of *M. incognita, Meloidogyne javanica*, and *Meloidogyne arenaria*, the three most widespread and agriculturally important MIG species. We also sequenced the genome of a second isolate of *M. floridensis*. Unlike most other MIG species, *M. floridensis* oocytes display some components of meiosis including chromosome pairing into bivalents [25]. As outgroup we sequenced *Meloidogyne enterolobii* (junior synonym *M. mayaguensis*), a highly pathogenic and invasive species, also apomictic. *M. enterolobii* is a member *Meloidogyne* Clade 1 like the MIG, but clearly distinguished from them by mitochondrial and ribosomal RNA sequence comparisons [20,26]. In this paper we examine the phylogenomic relationships between the species, investigate the origins of the apomictic species, examine hybridisation and ploidy, and describe levels of intra- and inter-specific variation. Together these comparative genomic approaches yield a detailed view of evolutionary history of these crop pests, and provide a valuable platform for improving future agricultural practices.

## Results

### Whole genome sequencing of nineteen isolates from five Meloidogyne species

*Meloidogyne* J2 larvae, egg masses, or bulk genomic DNA samples were obtained either frozen or preserved in ethanol. The samples represented isolates taken from diverse geographic locations and grown in laboratory culture before harvesting (Table S1, Supplementary Results). Isolates were identified to species by the source laboratories, identifications later confirmed by sequencing. High-coverage Illumina short-read data were produced from each isolate. The raw sequence data have been deposited with the international sequence databases under BioProject reference PRJNA340324.

### Reference genome assembly and annotation

We assembled one reference *de novo* genome for each of the five species, identified repeats and predicted genes on these assemblies, and then mapped data from other conspecific isolates to call variants. The isolates chosen for reference assembly (because of superior sample quality) were *M. incognita* W1, *M. javanica* VW4, *M. arenaria* HarA, *M. enterolobii* L30 and *M. floridensis* SJF1 (Table 1). We excluded contamination from our assemblies using the TAGC “blobplot” pipeline [27]. Very few primary assembly scaffolds (<1% of the total span) were removed as likely contaminants. We tested several genome assembly approaches, and found that Platanus [28] produced the most complete assemblies for the apomictic species, based on representation of core eukaryotic genes [29] and expected genome sizes (Table 1). For the automictic species *M. floridensis*, the Celera assembler [30] yielded the best assembly. Our assemblies are included in the archive associated with this publication (doi:10.5281/zenodo.399475).

**Table 1:** *Summary metrics for the reference MIG genomes*.

| Species | Strain or isolate code | Assembly Span (bp) | Number of scaffolds | Longest Scaffold (bp) | Scaffold N50 (bp) | CEGMA* |  | Number of predicted genes | %GC |
| --- | --- | --- | --- | --- | --- | --- | --- | --- | --- |
|  |  |  |  |  |  | C | A |  |  |
| *M. javanica* | VW4 | 142,608,877 | 34,394 | 223,460 | 14,133 | 90% | 2.5 | 26,917 | 30.2 |
| *M. incognita* | W1 | 122,043,328 | 33,735 | 248,829 | 16,498 | 83% | 2.4 | 24,714 | 30.6 |
| *M. arenaria* | HarA | 163,770,989 | 46,509 | 163,224 | 10,504 | 91% | 2.7 | 30,308 | 30.3 |
| *M. enterolobii* | L30 | 162,361,678 | 46,090 | 94,967 | 9,280 | 81% | 2.6 | 31,051 | 30.2 |
| *M. floridensis* | SJF1 | 74,893,904 | 9,134 | 88,393 | 13,256 | 84% | 1.6 | 14,144 | 30.2 |

\* CEGMA: Core Eukaryotic genes Mapping Approach [29]; C: % complete CEGMA orthologs; A: average copy number per CEGMA ortholog

We also sequenced the genome of *M. haplanaria* isolate SJH1 (BioProject PRJNA340324). *M. haplanaria* is outside of the MIG phylogenetically, and was only included in mitochondrial genome analyses (see below). We did not include the previously published *M. incognita* genome [9] in our analyses as it was produced with older sequencing technologies, and older assembly algorithms, and without access to the raw sequencing data we were unable to appropriately compare it (Supplementary Results, section 1).

Our reference genomes ranged in span from 75 Mb (*M. floridensis*) to 164 Mb (*M. arenaria*) (Table 1). The transposable element content of the genomes was characterised as described by Szitenberg et al. [31] (Figure S2, Supplementary Results). While estimates of the proportion of the genome occupied by mobile elements will be influenced by the accuracy of genome size estimates themselves, we found that Clade I RKN (MIG plus *M. enterolobii*) had a greater genome proportion of TEs than did *M. hapla*. However there were no clear differences in transposable element content between the MIG species or in relation to reproductive mode, as also discussed in Szitenberg et al [31]. We identified from 14,144 to 30,308 protein coding genes in the five reference genome assemblies (Table 1). The number of genes predicted in the *M. incognita* genome was higher than reported previously for this species (17,999 [9]), but for *M. floridensis* we predicted a similar number of protein coding genes to that reported previously (14,500 [10]). Protein coding gene sequences were recovered from other samples (listed in Table S1) using a mapping-and-assembly approach. Reads were mapped to the genes of the most closely related reference genome, based on their mitochondrial relationships (see below).

### Divergent genome copies are common in MIG genomes

Previous analyses of the *M. incognita* [9] and *M. floridensis* [10] genomes revealed that many loci were present as two divergent genomic copies, a pattern not observed in *M. hapla* [11]. These divergent copies have signatures of having been brought together by hybridisation [11]. Our analyses of all five of the newly assembled genomes, using a range of methods, indicated that divergent gene copies are a common feature of all MIG genomes (figures 1, 2 and 3) and not found in the Clade 2 automictic, diploid *M. hapla*. All the apomictic taxa show a leptokurtic peak of within-genome, non-self best BLAST matches with a mode at ~97% identity (Figure 1). Notably while *M. floridensis* also had a peak of pairs at ~97% identity, there were many fewer such loci in both our newly assembled *M. floridensis* SJF1 genome and the previous assembly [10]. To robustly define groups of orthologous genes across our species set, we clustered the protein sequences from the five reference genomes with OrthoFinder [32]. We found that the default inflation parameter (1.5) merged what appeared to be distinct sets of orthologs, and so used a more restrictive inflation value of 2. A total of 29,315 orthology groups (OGs) were recovered. Within these we selected the 4,675 OGs that contained from 1 to 4 gene copies in all *Meloidogyne* species. We filtered the OG predictions to remove likely artifacts of over-clustering, and recent within-species duplications. We identified recent paralogs (in-paralogs; where sequences from the same species were close sisters) and retained only the less divergent of the two gene copies. To avoid incorporation of groups that contained more than one set of orthologs, we verified that the sequences from *M. enterolobii* were monophyletic. This was true of all OGs that had passed the first filter. This filtered dataset contained 3,544 OGs, with one or two orthologs per species in most groups (Figure 2). Although a third divergent genomic copy appeared to be present in for a few loci, reanalyses of these data revealed that these were largely likely to be derived from in-paralogs or fragmented gene predictions (Supplementary Results, section 2). For the 3 apomictic species, there were 1632–1920 orthology groups containing two copies, but many fewer were seen for *M. floridensis* (477), consistent with the intragenomic analysis (Figure 1). Many groups contained two gene copies in more than one species, with 871–1046 shared between apomictic species pairs and 225–246 shared between *M. floridensis* and the apomictic species. We selected a subset of 612 OGs which contained two copies in at least three of the five RKN species. We removed OGs containing possible gene prediction artifacts (i.e. where a species was represented by putative homoeologs with less than 20 bp overlap, as these may have derived from single, fragmented gene copy). While a few OGs included three gene copies from individual MIG species, the proportion of these “triples” was lower than reported previously [9,10]. These filters removed a further 75 OGs, leaving 533 OGs for further analysis.

**Figure 1:**
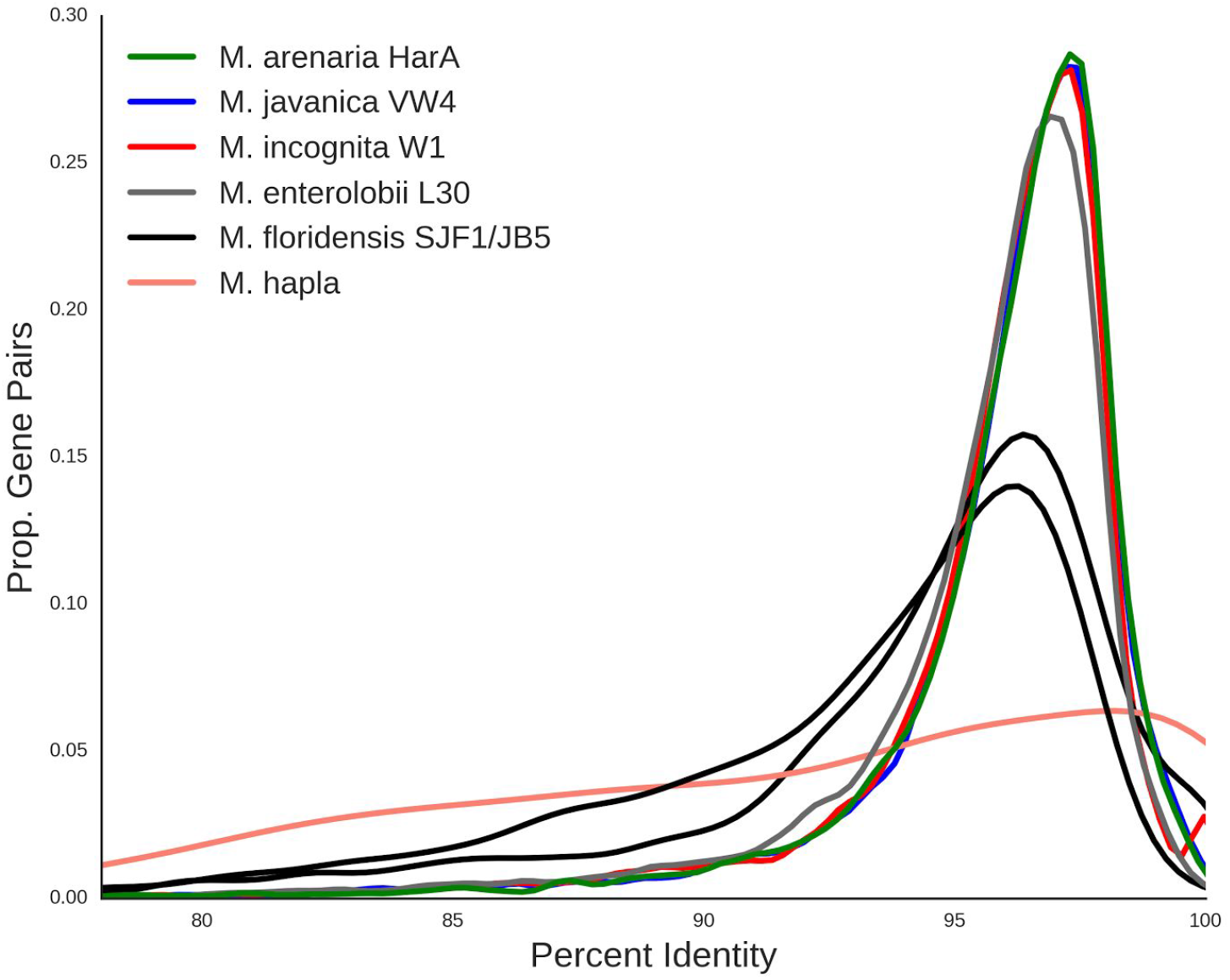
Meloidogyne incognita-group nematode genomes contain many gene pairs with similar sequence divergence. For each species, we identified the next-best BLAST hit for each gene in its source genome, and calculated the pairwise identity. This smoothed density plot shows the the proportion of gene pairs at different percent sequence identity for each species analysed. The x-axis is the percent identity between gene pairs and the y-axis is the proportion of gene pairs with a given percent identity. The histogram was smoothed by kernel density estimation.

**Figure 2:**
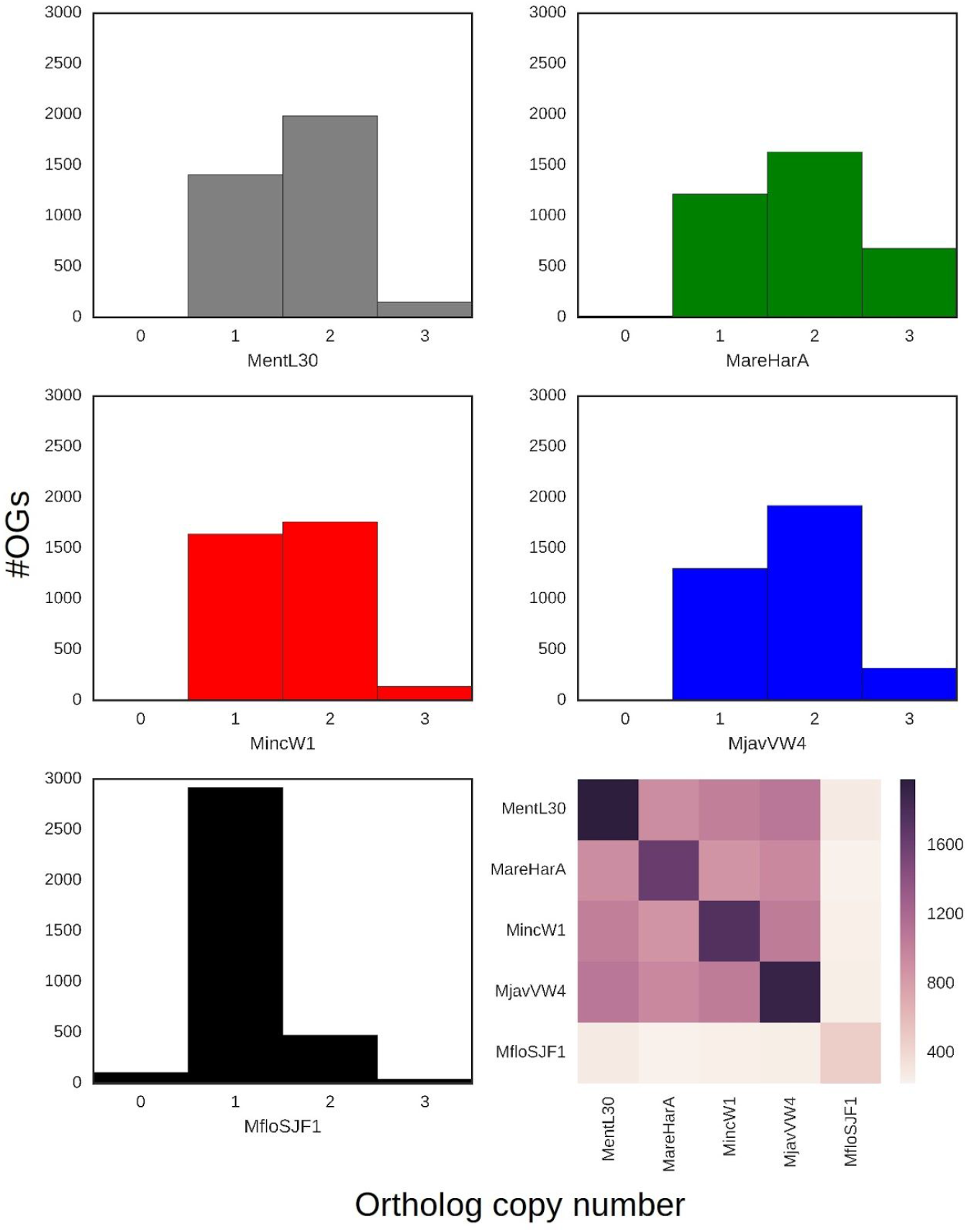
Ortholog counts in *Meloidogyne* genomes. Histograms of ortholog copy number within orthology groups (OG) for each reference genome assembly (panels 1–5). The heatmap (panel 6) represents the number of OGs containing two copies found in a genome on the diagonal, and how many of those copies are shared between species. Figure S4 (Supplementary Results) shows the number of shared OGs with one and three copies.

**Figure 3.**
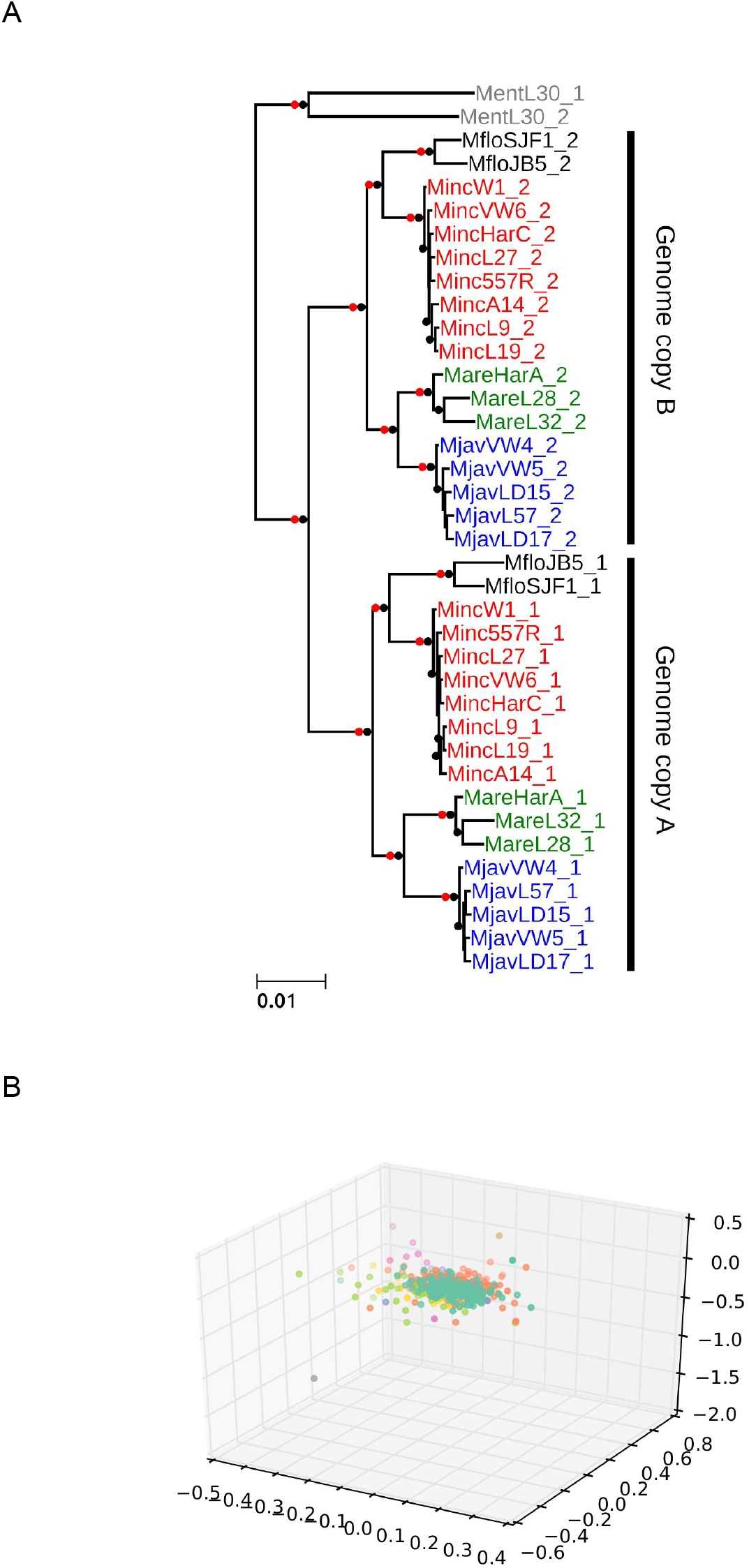
Phylogenetic analysis of gene sets in MIG species. A: A maximum likelihood phylogenetic tree based on 533 nuclear loci reveals that the separation of “A” and “B” gene sets is internally congruent (*i.e*. all of the pairs are likely to have originated at the same time). This event predates speciation of the MIG taxa. An independent phylogenetic origin is shown for the divergent gene pairs in *M. enterolobii*. Nodes with maximal bootstrap support are denoted by black bullets. Nodes that were recovered in the homoeolog randomization analyses (Supplementary Results, Figures S5 and S6) are marked by red dots. The topology is very similar to the one recovered from a coalescent phylogenomic approach (Supplementary Results, Figure S5B) and, for each of A and B subtrees, also similar to the one recovered for the mitochondrial genome phylogeny (Supplementary Results, Figure S7). B: Concordance in gene trees supporting a single-origin scenario. Gene tree cluster analysis was used to test for conflicting phylogenetic signal in the phylogenomic dataset. Maximum likelihood gene trees were reconstructed for each of the 533 orthology groups (see Methods for the determination of orthology and identification of homoeologs). A pairwise weighted Robinson-Foulds distance matrix was computed between all gene tree pairs. Only a single cluster was recovered when embedding the distance matrix in 3D space via metric MDS. Attempts to enforce up to 10 distinct groups (colours) did not reveal any separation.

### Phylogenomics of MIG species and genomes

We collated sequences corresponding to the set of 533 OGs in which two gene copies existed in at least three of the MIG species from the nineteen whole genome sequenced isolates. We generated a supermatrix alignment where each MIG isolate and *M. enterolobii* were represented by two operational taxonomic units (OTUs), “A” and “B”. We randomly assigned the paired gene copies from each species to the corresponding “A” or “B” OTU. Maximum likelihood (ML) phylogenetic analysis yielded a well resolved tree with the MIG “A” and “B” OTUs as sister clades (Figure 3A). Within each of “A” and “B” the several isolates of each species were robustly grouped together, and the branching order of these species was identical in the “A” and “B” subtrees, splitting the four taxa analysed into two groups of two: *M. floridensis* with *M. incognita*, and *M. javanica* with *M. arenaria*. Based on the sequenced isolates, divergence within species between isolates was very low for *M. incognita* and *M. javanica*, and slightly larger for *M. floridensis* and *M. arenaria* (Figure 4A). The phylogenetic relationships between species for each genome copy were supported with maximal bootstrap support (black nodes in Figure 3A). We conducted two randomization analyses in which for each OG we shuffled homoeolog identity between OTUs “A” and “B” and reconstructed trees based on ML analysis of randomized supermatrices or on coalescent analysis of randomized gene trees (see Methods, Supplementary Results Figures S5 and S6). Both randomisation tests supported all the same nodes (red nodes in Figure 3A). Coalescent phylogenomic analyses yielded the same topology (Supplementary Results, Figure S5). The two *M. enterolobii* OTUs were resolved as monophyletic, outside the MIG clade, indicating that the divergent gene copies in *M. enterolobii* had an evolutionary origin distinct from the event that generated the MIG genome copies.

**Figure 4.**
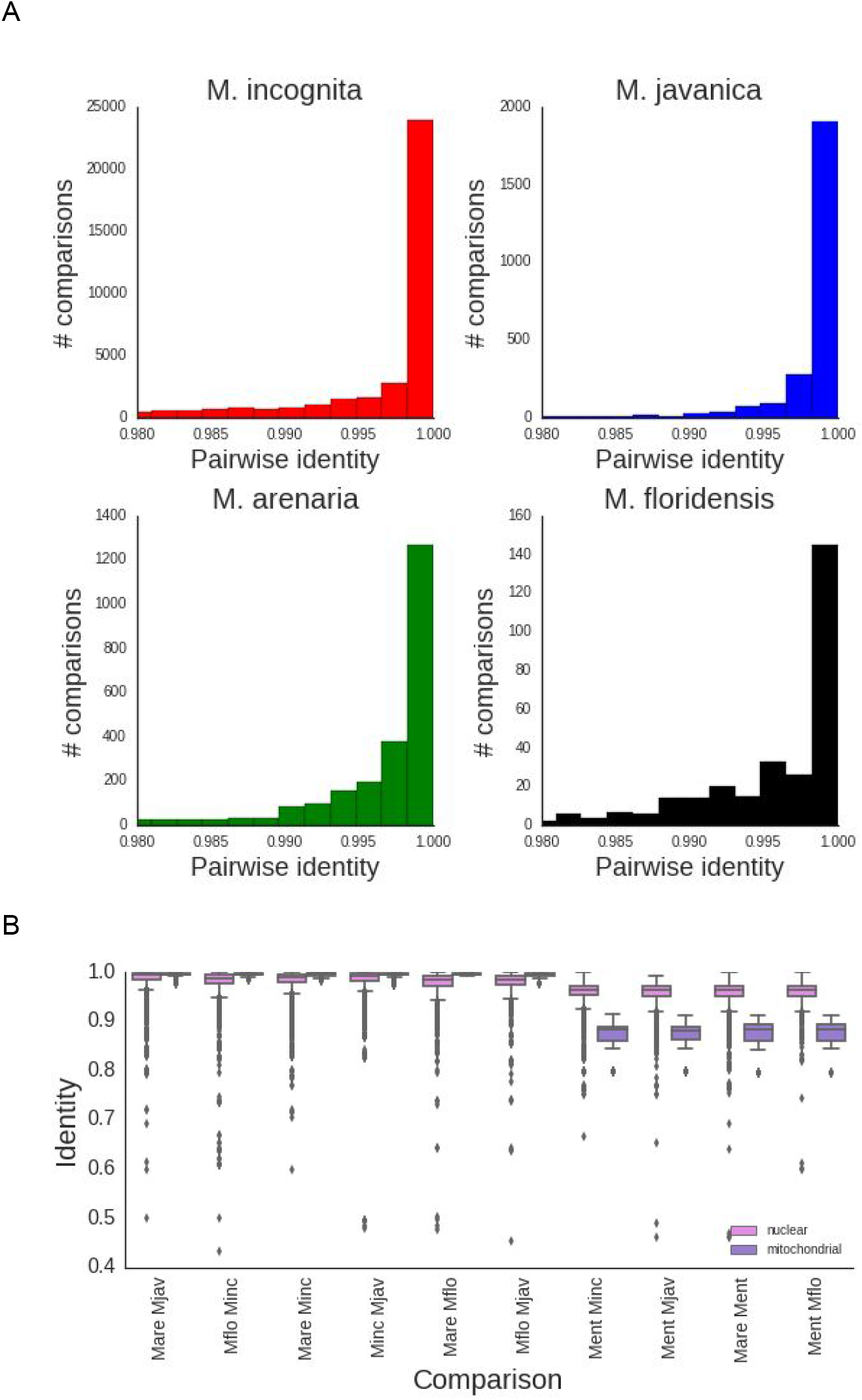
Nuclear and mitochondrial sequence variation in *Meloidogyne*. A: Pairwise identity of nuclear protein coding genes between isolates of each MIG species. B: Pairwise identity of nuclear and mitochondrial genes between MIG species and between MIG and outgroup RKN species. The rRNA loci and the putative Control Region were included in the mitochondrial dataset. Mare: *M. arenaria*; Mjav: *M. javanica*; Mflo: *M. floridensis*; Minc: *M. incognita*; Ment: *M. enterolobii*.

Simultaneous analysis of many loci in a phylogenomic reconstruction can obscure the presence of alternative phylogenetic histories for subsets of those loci. In order to test for this within our dataset, ML trees were constructed for each orthology cluster and a pairwise weighted Robinson-Foulds distance matrix was computed between all gene tree pairs. Only a single cluster was recovered when embedding the distance matrix in 3D space via metric MDS (Figure 3B). Attempts to highlight up to 10 distinct groups did not reveal any separation, supporting the existence of a single topology.

Mitochondrial coding sequences showed little divergence within and among MIG species (median identity across genes > 99%, Figure 4B), and we were only able to resolve species relationships with complete mitochondrial genome or complete mitochondrial coding sequence datasets. Even with the mitochondrial genome dataset there were just 78 out of 10,955 phylogenetically informative sites. The MIG were quite distant from the outgroups, with *M. haplanaria* closer than *M. enterolobii*. Despite the lack of strong phylogenetic signal, the mitochondrial phylogenies were congruent with the nuclear genome analyses, with the exception of the paraphyly of *M. arenaria* (Supplementary Results, Figure S7).

Surprisingly, intraspecific sequence identity was lower in the nuclear genome (median values of 0.987–0.994; Figure 4A) than in the mitochondrial genome (identities were 1.0 with the exception of *M. arenaria*, where the median identity is 0.997; Figure 4B). *M. incognita* and *M. javanica* had lower intraspecific diversity than did *M. arenaria* and *M. floridensis* (Figure 4A), even though fewer isolates were examined for the latter two species, and in accordance with branch lengths in Figure 3A.

### Evidence of genetic exchange between homoeologs

Above we showed that in the MIG apomict species there is strong evidence for large-scale presence of two distinct copies of many nuclear genes, and multiple analyses of the evolutionary histories of these copies were congruent. These findings suggest that the MIG species have two distinct genome copies generated by the same, major genome event and that these genomes came together or existed in an ancestor of the four MIG taxa analysed. This event could explain the increased assembly span and elevated protein coding gene number observed in these species compared to the homozygous diploid species *M. hapla*. However, not all genes in the MIG taxa were present in divergent copies (Figure 1). Also, while more genes in the apomict MIG taxa *M. incognita, M. javanica* and *M. arenaria* were present in the assembly in divergent copies than as a single copy, in the automict *M. floridensis* only a small proportion of the OGs contained two divergent copies (Figures 1 and 2).

If the MIG species’ genomes are the product of a process that resulted in, originally, two divergent copies of every gene, then loss of one copy of a subset of genes would need to have occurred. Stochastic deletion of second copies in a piecemeal fashion is one possible explanation. However this appears *a priori* highly unlikely for the cytogenetically diploid *M. floridensis*. Another mechanism would be ameiotic non-crossover recombination between homoeologous chromosomes resulting in two homozygote copies on one side of the recombination event while heterozygosity is maintained on the other side (apparent gene conversion), or double strand break repair mechanism also resulting in gene conversion.

To identify and quantify such potential non-crossover recombination events, we used a sliding window BLAST approach within and between species (see Methods and Supplementary Results, Figure S8A). We identified locations in a query genome in which two overlapping windows in the query had the same top two best hits in the target genome, except that the best hit in the first window was the second best in the second, and *vice versa*. Each pair of such overlapping windows was counted as an event. To distinguish gene conversion from reciprocal crossover, we checked that exchanging the query and the target did not yield the same locus. In the apomict species all the detected events had the signature of non-crossover recombination as we did not detect reciprocal crossover. Gene conversion usually results in a short conversion tract (less than 100 bp).

While we observed such short conversion tracts by manual inspection, further work is required to quantify this subtle signature throughout the genome and to determine the extent of its contribution to the observed genetic exchange between homoeologs.

For the meiotic *M. floridensis*, our assembly span was shorter than would be expected from a species carrying two distinct genome copies (Table 1) and the genome appeared largely homozygous. A homozygous genome does not provide the contrast to detect crossover or recombination because such events will have no consequence on the sequence of either the identical genome copies. Within the apomicts, non-crossover recombination rates are largely explained by phylogenetic distances (Pearson’s r = 0.73), suggesting a steady rate of such recombination, although when *M. floridensis* is included this relationship weakens (r = 0.4) (Supplementary Results, Figure S8B).

### Read coverage analyses and estimation of ploidy

Based on cytological examination, chromosomes in MIG apomicts are very small and their number varies within and between species (Triantaphyllou 1981; 1985). This chromosome number variation has led to the speculation that many of these species/isolates are aneuploid (hypotriploid). A previous assembly of the *M. incognita* genome also predicted some level of triploidy [9] but some doubt exists regarding the evidence presented there (Supplementary Results, section 2). We assessed the likely ploidy represented by our assemblies through read coverage analysis, and the stoichiometry of the divergent gene copies. We identified ~350 contig pairs, shared among the three apomictic MIG species, that had shared content of diverged gene pairs (only ~100 contig pairs were shared between the MIG apomicts and *M. enterolobii*). We normalised the modal coverages for each contig by expressing them as a ratio of “alpha” to “beta”, where the copy with greater coverage was arbitrarily designated “alpha”. Thus a coverage ratio of 1 indicates one copy each of alpha and beta, while a ratio of 2 would indicate two copies of each alpha segment to each beta segment, or three copies overall.

In all species a bimodal Gaussian distribution fit the data better than did a unimodal Gaussian distribution (15 < Δ Residual error < 50) (Figure 5). This result conflicts with the presence of a simple stoichiometry (one copy each of alpha and beta) across the whole of the genomes of these apomicts, as some genomic regions fit a triploid model (alpha1, alpha2, beta). The alpha1 and alpha2 genomic copies are almost identical in sequence within a genome. Many of the same genomic regions are present in double stoichiometry across the MIG genomes (Figure S11), suggesting that they were largely formed before the MIG apomict speciation.

**Figure 5.**
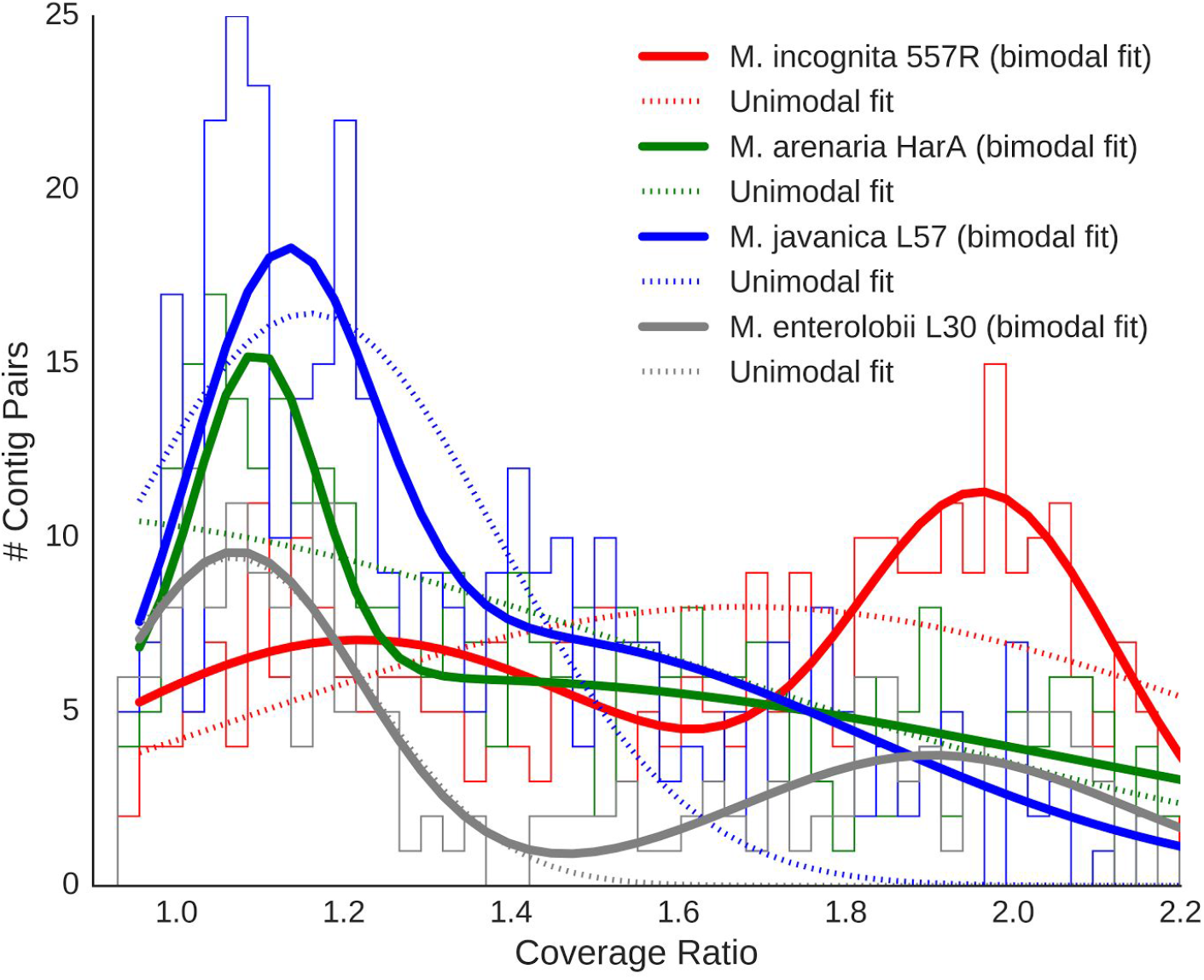
Coverage data suggest hypotriploidy in MIG genomes. Histograms of relative coverages of A and B genome contigs in several MIG genomes and *M. enterolobii*. The solid lines represent binomial Gaussian and the dotted lines unimodal Gaussian models of the data.

**Figure 6:**
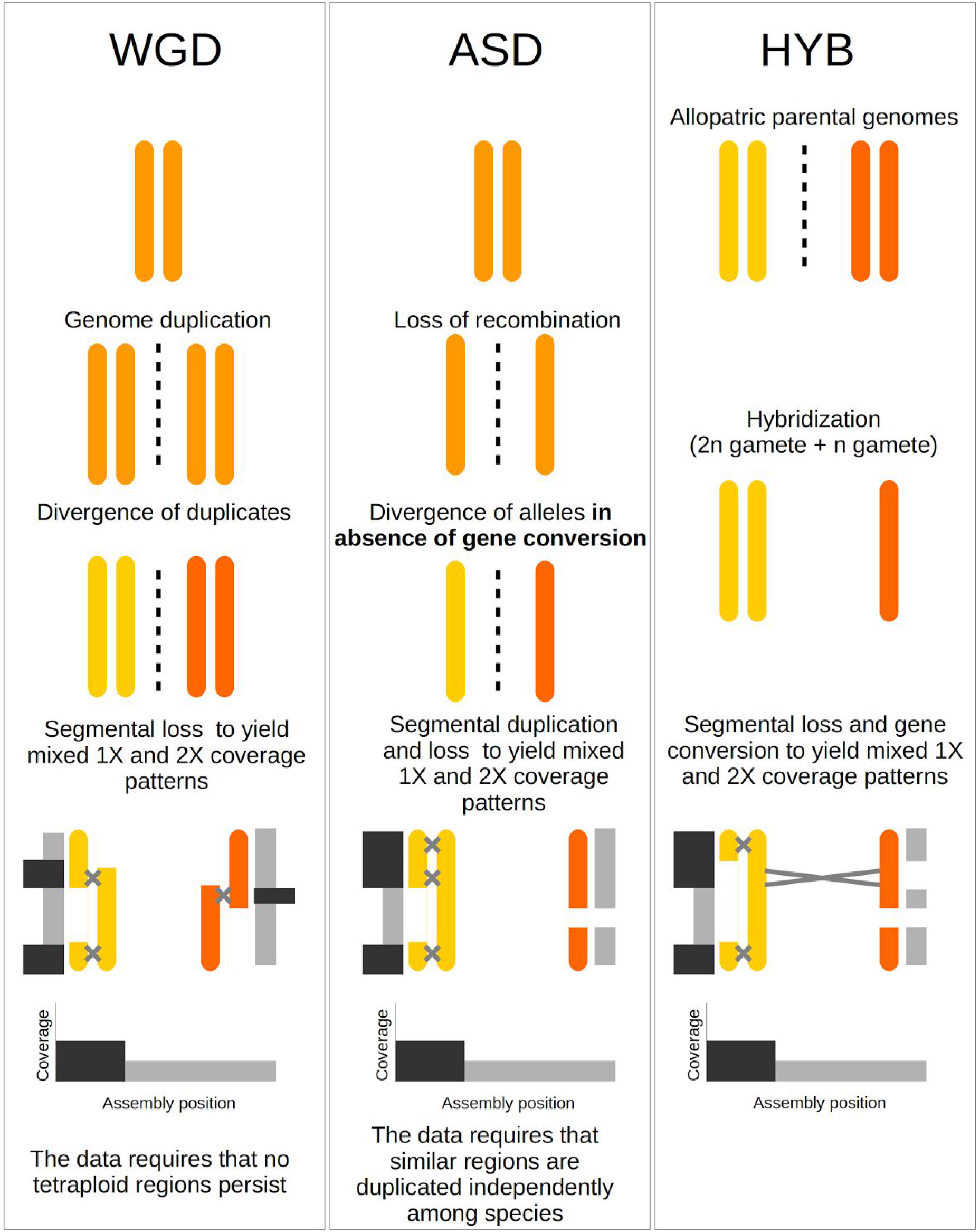
A model for the hypotriploid MIG genome. Three scenarios are considered, including whole genome duplication (WGD), frozen allelic sequence divergence (ASD) and hybridization (HYB). HYB is the most parsimonious scenario given the phylogenetic relationships of genomes A and B (Fig 3), because it does not require the complete loss of tetraploidy (WGD) or independent but identical duplication events across species (ASD). In all scenarios, speciation would have occurred just before or during the last step. Grey and black bars parallel to the recombining chromosomes represent their read depth in the assembly, with black bars representing twice the coverage than grey bars. All the scenarios can produce similar coverage values across the assembly.

The relative proportions of the genome assessed as being present in 1:1 stoichiometry *versus* 2:1 stoichiometry differed among species (Figure 5). Median coverage ratio distributions for all the isolates (Supplementary Results Figure S9 and summarized for species in Figure S10) showed that *M. incognita* had the highest fraction of the genome in the 2:1 pattern, and *M. enterolobii* the lowest. The proportion of the loci present as homoeologs in *M. enterolobii* and in the MIG apomicts was similar (Figure 1), but different orthology clusters were represented in the duplicated fraction of *M. enterolobii*. While more than 350 pairs of contigs carrying divergent copies among the MIG apomicts, only 112 of these genome segments were shared between the MIG apomicts and *M. enterolobii*. This finding supports the hypothesis, derived from phylogenetic analyses (Figure 3A), that the MIG and *M. enterolobii* represent independent origins of divergent genome copies.

## Discussion

We have generated good-quality assemblies of the genomes of five agriculturally damaging species of plant-parasitic root-knot nematodes. We also generated whole genome resequencing data for several isolates of the apomictic MIG species, and have analysed our data for insights into the processes that have generated their peculiar genome structures, in the resulting patterns.

We have presented four main findings. Firstly, the assemblies of the apomict MIG species (Table 1) have spans and gene counts much greater than observed in the diploid *M. hapla*. Secondly, all MIG genomes harbour a substantial number of pairs of divergent gene copies (Figure 2). Similar divergent gene copies are absent from the diploid species *M. hapla*, and are reduced in number in the meiotic MIG species *M. floridensis*. Phylogenetic analyses of these divergent gene pairs (Figure 3) showed that they have congruent evolutionary histories, and that they likely arose from a single, major event before the speciation of the four MIG taxa analysed.

Thirdly, analysis of read coverage of the apomictic genomes revealed that many of the duplicated segments had a 2:1 stoichiometry (Figure 5). Coupled with historical karyotypic analysis of MIG species, which has revealed greater than diploid chromosome numbers, these data are best reconciled with partial triploidy of the MIG species, with two copies of one genome and a single copy of the other. The two copies of one genome are very closely related (most genes are identical, and maximal divergence was 0.1%), whereas the other, divergent genome is approximately 3% different in coding regions and much more divergent in noncoding regions.

Lastly, we found that not all genes were present as divergent duplicate copies, suggesting that there has been stochastic loss of heterozygosity in some divergent gene copies. Homozygous genes were frequently shared between MIG species, implying that these events have been ongoing through speciation. Exploring them we found evidence for frequent recombination events, most likely non-crossover events, where discordant similarities mapped along assembly scaffolds showed replacement of one copy by its sister.

### MIG species possess divergent *A* and *B* genomes

From these data we suggest that the MIG species have complex, hypotriploid genomes, made up of divergent *A* and *B* homoeologous subgenomes. Arbitrarily, we designate the effectively diploid subgenome as *A* (and thus *A1* and *A2* copies), and the other genome as *B*.

The phylogenomic tree was well supported, and the data displayed no detectable conflicting signal (Figure 3). Randomly partitioning gene pairs into “A” and “B” sets generated congruent species phylogenies for each set (Figure 3). Genome assembly quality was not an issue, as the two isolates of *M. floridensis* grouped together robustly despite the more fragmented first *M. floridensis* draft genome [10]. Both of the divergent copies yield the same species phylogeny, and thus the uniting of the *A* and *B* lineages in a common progenitor must have occurred before the speciation of the MIG. *M. enterolobii* is well separated from the MIG species (Figure 3), as has been reported previously [18,21,33]. *M. enterolobii* is also an apomict, and also contains divergent genome copies, but the phylogeny show that these copies arose as a different progenitor from the one at the base of the MIG. *M. enterolobii* thus likely represents an independent origin of apomixis. Transitions to apomixis have happened on at least four separate occasions elsewhere in the genus *Meloidogyne* [7]. However, the relationship of cause and effect between the emergence of divergent genomes and the loss of meiosis remains to be deciphered.

### The origins of MIG A and B genomes

We do not known what mechanisms led to the presence of the two divergent genomes in the MIG species. Also it is not known when in their evolutionary history apomixis occurred. Below we propose three broad mechanisms to account for the divergent *A* and *B* genomes found in the MIG.

#### Whole genome duplication (WGD)

Whole genome duplication in the ancestor of the MIG would create a tetraploid, then driven to hypotriploidy by gene loss, but with no evidence of tetraploid regions in the MIG genomes, it is unlikely. Additionally, duplicated genomes would accumulate mutations before the MIG species began to diverge as the *A* and *B* genomes differ by ~3% in protein coding regions, but sequence divergence among species is much smaller. However, we find that non-crossover recombination largely homogenizes the *A1* and *A2* genomes found in triploid regions and also some homoeolog regions of the *A* and *B* genomes. Such a differential homogenizing effect required to justify WGD in MIG is unparsimonious.

#### Frozen allelic sequence divergence (ASD)

A transition to apomixis could instantaneously ‘freeze’ the original haploid chromosomes (alleles) of the parent as independently evolving entities. Organismal levels of ASD vary greatly across different taxa [34] and apomictic species can have very high ASD, due to lack of recombination. The resulting sub-genome divergence under this model could be very similar to that of a hybrid genome. However, hypotriploidy is again not easy to explain also in a frozen ASD model. The loci present in a triploid *alpha1, alpha2, beta* arrangement are largely shared by the MIG (Figure S11) such that they most likely emerged in a shared ancestral event. ASD in MIG apomicts, coupled with hypodiploidy, could only be explained by invoking extensive segmental duplication before species divergence, followed by suppressed duplication since speciation, which also seems unlikely.

#### Hybridisation

Hybridisation between two diverse RKN strains followed by loss of meiosis is an attractive explanation for the genomic structure of MIG species since it accounts for all our observations in a single step. Hybridisation allows contributing genomes to have diverged in allopatry (separate organisms, species, or regions) so that recombination of any kind did not act to homogenise the *alpha* and *beta* alleles. Upon hybridisation genes are pre-diverged, and can be considered homoeologs. Hypotriploidy can be readily explained by an interspecific hybridisation event involving one reduced (n) and one unreduced (2n) gamete followed by a rapid genomic turnover back towards diploidy. This is a process well described in the literature, especially concerning interspecific hybridisation [35]. We note that hybrid origins for intra-genomic divergent alleles are very well documented in both animals and plants [36–40], whereas extreme ASD by asexual accumulation of mutations is controversial, with few if any clear examples. The bringing together of diverged chromosomes may be a mechanism contributing to the disruption of meiotic segregation and thus the origins of asexual modes of reproduction, and Janssen et al [7] indicate an association between cytological triploidy and apomixis in *Meloidogyne*.

We note that not all loci are present as divergent *alpha* and *beta* gene pairs, and the proportion of the genome with elevated coverage does not correspond to half of each assembly. The MIG species differ in their proportions of loci present as divergent pairs, indicative of independent change in each lineage. *M. incognita* had approximately half the loci as divergent pairs (Figure 2).

The predictions of frozen ASD and hybridisation models do diverge for the source populations. Hybridisation predicts that divergent parental species exist (or existed) and that each have (had) genomes more related to one of the MIG homoeologous genomes than to the other. These parental lineages may not be common in agricultural environments and have not been recorded. The discovery of wild species with non-hybrid genomes, which were phylogenetic sister to either the A or B MIG genomes, would be key to our understanding of MIG origins.

Another characteristic of MIG species that likely contributes to their genome complement and evolution is the chromosome structure of this group. As is true throughout the phylum Nematoda, *Meloidogyne* chromosomes are holocentric. In addition, based on cytological examination, chromosomes in MIG apomicts are very small and their number varies within and between species (Triantaphyllou 1981; 1985). For most isolates of *M. incognita*, the chromosome numbers are between 41 and 48 and they are considered to be hypotryploid. Considerable polymorphism between isolates has also been observed for relative size distribution of chromosomes (Triantaphyllou, 1981). These features suggest that chromosome missegregation, breakage, translocation, inversions or similar events may have contributed to the genome copy variations seen in the modern MIG species. Cytogenetic studies using modern techniques would be interesting and necessary to explore these possibilities.

### *M. floridensis* is a sibling not a parent of *M. incognita*

In contrast to the ameiotic species, *M. floridensis* appears to be effectively diploid from both cytological [25] and genomic analyses (this study). We show (Figure 2) that the process of homoeolog loss is almost complete in *M. floridensis*, with the telling exception of the gene pairs we use in the phylogeny to reveal its more complex history. Despite sharing the same hybrid origins (Figure 3A) *M. floridensis* has regained or maintained the ability to carry out meiosis in a form of automixis. How it was able to maintain a meiotic reproductive system, during this turbulent and dynamic period of genome reorganisation, and whether the more complete re-diploidisation is a cause or a consequence of the retention of meiosis, will require further study.

Lunt et al [10] hypothesised, based on shared homoeologous genes and difference in reproductive mode, that *M. floridensis* was a parent of the hybrid MIG apomicts. Our phylogenomic analyses, using much richer data, rejects this hypothesis. Instead *M. floridensis* is a close relative of *M. incognita*, is derived from the same parents, and nested within the apomict MIG species. Handoo et al [25] showed that *M. floridensis* is cytologically very different from the MIG apomicts, and enters meiosis during oogenesis. Eighteen chromosome pairs were observed, and bivalents were present before nuclear division and polar body formation, though no second meiotic division was observed. This strongly suggests that meiotic recombination occurs in *M. floridensis*, as is usual in automixis. The Clade 2 RKN *M. hapla* also reproduces by automixis [41] and in this species meiotic parthenogenesis results in rapid homozygosity (as meiosis II is not completed and diploidy is restored by rejoining of sister chromosomes). If *M. floridensis*’ meiosis involves endo-reduplication and failure to complete meiosis II as suggested by Handoo et al [25] this would explain the substantial loss of heterozygosity compared to its closest relative, *M. incognita*.

Additionally, Lunt et al [10] proposed a complex double-hybridisation to explain the relationship between the *M. floridensis* genome and the published *M. incognita* genome [9]. Sections of the *M. incognita* genome were reported to be present in three diverged copies [9], implying that three separate parental genomes had been brought together from, we inferred, two hybridisation events. There have been many suggestions in the literature that *M. incognita* is triploid or hypotriploid, based on cytological data, and the observation of triploidy indicated by three diverged copies of some loci by Abad et al [9] was biologically plausible. We also found some instances of three diverged copies in our initial assemblies, and *M. arenaria* has several loci present as three diverged copies (Figure 2). However careful re-examination of these (Supplementary Results) revealed that they were all gene prediction artefacts, problematic orthology groups (merging two genes, or representing one fragmented ortholog as two separate copies), or paralogous gene family members. We find no convincing evidence for the presence of three homoeologs in any of our sequenced MIG genomes, or in our re-analyses of the published *M. incognita* or *M. floridensis* genomes. Although we discuss above that MIG species do indeed have hypotriploid genomes, none contains a third divergent genome copy, but rather a second copy of one of the two homoeologs found in all other MIG species.

### Genome size and gene number in the MIG

The genome sizes and gene numbers in the MIG species do not fully correlate with predictions of hypotriploidy. Many current genome assemblies come from organisms that are inbred, or naturally homozygous, leading to alleles being merged in the genome assembly, and the production of a collapsed, haploid assembly from a diploid organism. Assembly of the genomes of organisms that have greater divergence between alleles may result in a partially uncollapsed, near-diploid genome estimate, where some of the genome generates a collapsed haploid sequence, while divergent segments are independently assembled. If we assume, as seems likely from our analyses (Figure 2), that homoeologs derived from *A* and *B* genomes are present in the apomicts for about half of the genomic regions, and that they are represented separately in the assembly, then our reference genomes (Table 1) will represent partially diploid assemblies, and be expected to be larger than a comparable haploid assembly. The 53 Mb genome assembly of *M. hapla* is high-quality and contiguous, and matches closely to experimental estimates of the haploid genome size (50 Mb) [11]. The sequenced *M. hapla* was highly homozygous, and the assembly is expected to contain no or very few regions where haploid segments have assembled independently due to sequence divergence. If *M. hapla* represents a base genome size for *Meloidogyne*, we would expect the assemblies from the MIG apomicts to be of the order of 75 Mb. However these species generate assemblies that are 122 to 163 Mb. Genome size is a biological attribute and varies between even closely related species, due to segmental duplications, expansions of repetitive elements and transposons, and long-term biases in insertion versus deletion. Szitenberg et al [31] showed that the MIG have an increased content of transposable elements compared to *M. hapla* (Supplementary Material, Figure S1). The larger MIG apomict genome assemblies are thus likely a reflection of this biological variation in genome size. The *M. floridensis* assembly is ~75 Mb in span but largely homozygous, and is, as expected, between *M. hapla* and the MIG apomicts in size.

Protein coding gene number estimates are influenced by annotation procedure and genome contiguity in addition to real, biological, sources of variation. Poor genome quality can lead to gene number inflation by fragmentation of predicted coding sequences, or gene number reduction by poor assembly. We found between 14,144 protein coding sequences in *M. floridensis* and up to 30,308 (*M. arenaria*) in the apomicts (Table 1). The homozygous genome of *M. hapla* contains approximately 14,700 CDS [11] very similar to the number predicted in the mostly homozygous *M. floridensis*. The elevated gene numbers in the MIG apomicts likely reflect the independent prediction of genes in the homoeologous segments.

### Homoeolog loss and the evolution of MIG genomes

Gene conversion is an outcome of non-crossover recombination between chromosomes and is a common and important force influencing genome structure and diversity [42–44]. Gene conversion is associated with double strand break repair and involves the replacement of one, typically allelic, sequence by another such that they become identical. It is common during meiosis, which is initiated by a double strand break, but additionally occurs during mitosis [42]. We found multiple lines of evidence suggesting that this may be an important process in shaping MIG genomes.

We have provided evidence that a part of the genome of the apomict *Meloidogyne* is triploid containing *alpha1, alpha2* and *beta* copies of homoeologous segments. The apomict *alpha1* and *alpha2* regions contain the same protein coding loci in different species (Supplementary Material Figure S11), strongly suggesting that they arose in a common event in the MIG ancestor. The alternative, that three independent recent duplications occurred involving exactly the same genomic regions, is a less parsimonious scenario. *Alpha1* and *alpha2* are almost identical in sequence and they do not independently assemble, map, or resolve phylogenetically to indicate sequence divergence. This is challenging for the evolutionary scenario shown in the phylogeny (Figure 3A). If the *alpha1* and *alpha2* copies derive from the two haploid copies present in the *alpha* parental species, and have been evolving independently since that event, then *alpha1* and *alpha2* should be at least as divergent as the *alpha* genomes in different MIG species. The MIG species are clearly distinct with ~2% CDS divergence between them in each genome copy, and yet within a species the *alpha1* and *alpha2* copies have remained essentially identical over the same time span. The homogenisation of sequence copies is known in other genomes however. Concerted evolution is a type of gene conversion, found in most organisms, which operates repeatedly over large genomic regions [45] to maintain their sequence identity. It has been well-characterised in many systems including the homogenisation of eukaryotic rRNA repeats [46] and the palindromes of the male specific region on human and chimp Y chromosomes [47]. Concerted evolution will be more frequent between highly similar sequences, such as the original *alpha1* and *alpha2* alleles, than between more divergent sequences, such as *alpha* and *beta* homoeologs [42], which is a scenario supported by our own data.

Different apomict species have different proportions of *alpha* and *beta* homoeologs remaining in their genomes (Figure 2). One mechanism by which this could occur is the deletion of one of the divergent genomic copies, and deletions to restore diploidy are known in other polyploids [48,49]. Another mechanism by which homoeologs could be lost, but without deletion of any genomic region, is gene conversion. This process could homogenise A and B genomic copies, leading to a reduction of gene clusters containing two copies (Figure 2) with no way to regain these divergent sequences in an apomictic species. This process would likely be stochastic, occurring differently in each species lineage, and generating variability between the MIG apomicts. These outcomes are observed and in addition we see sequence homogenization. When the sequence of homologous contigs containing divergent A and B gene pairs are analysed we can directly observe the action of non-crossover recombination between those sequences.

Although converted sequences are homozygous, non-crossover recombination can be a mechanism that also increases diversity among mitotically reproducing organisms. For example, the hybrid plant-pathogenic oomycete *Phytophthora sojae* has a high rate of mitotic gene conversion between homoeologous genes, and different lineages of the pathogen have generated variation in avirulence by gene conversion of different genomic regions [50].

Although *M. floridensis* possessed the same A and B genomes, with the same evolutionary history, its genome is very different in its homoeolog content to that of the apomicts. Only a minor component of the *M. floridensis* genome is still present in divergent *alpha* and *beta* homoeologous copies (Figure 2) despite the fact that it shares genomic origins with the other MIG. The reduced genome span and count of protein coding loci in *M. floridensis* compared to the apomict MIG (Table 1) could also be a product of co-assembly of homogenised homoeologous loci. Extensive non-crossover recombination may provide a mechanism to explain these results. Since gene conversion is initiated by double strand breaks, which are associated with meiotic recombination, the automict *M. floridensis* may have an increased rate of gene conversion compared to the apomicts. This is very challenging to study since the loss of homoeologs itself removes the variation necessary to measure this process, and will likely require more data from different *M. floridensis* lineages. Meiosis is a diverse and powerful process that provides other mechanisms than gene conversion however by which to homogenise homologous sequences. *Meloidogyne hapla*, also an automict, homogenises its genome rapidly by the rejoining of sister chromosomes during meiosis [41]. Although superficially similar to the automixis in M. *hapla* the exact nature of the meiotic division of *M. floridensis* is still unclear [25]. If, as suggested, endoduplication of the genome is involved then this could also be a mechanism for homogenisation and loss of homoeologs not requiring high levels of gene conversion. Genetic pedigree and cytological studies will be informative in determining the exact nature of automixis in *M. floridensis*, but at present we cannot distinguish the driving forces for its genome-wide homoeolog loss.

Whether the differences in retention of ancestral homoeologous variants may underlie differences in host range and pathogenicity between species is a challenging question. The genome-wide differential retention of A and B genomes, as well as the presence of A1 and A2 copies, could be functionally very significant for these nematodes. The complexity of MIG genome structure, as well as the diverse genetic mechanisms that may be driving this, could be an important source of adaptive variation. These processes could operate to generate diversity between the asexual mitotic parthenogen species, but also by putting the meiotic parthenogen *M. floridensis* on a separate genomic trajectory. Understanding the nature and rate of adaptive evolution in MIG species, as they compete with the defences produced by plant resistance genes, will be an important direction for future research.

### Low intraspecific divergence implies recent global colonisation

The most economically important MIG species are globally distributed in agricultural land across the tropics, and whether they are recent immigrants or more anciently endemic in those locations has been unclear [2]. Comparisons of mitochondrial genomes have revealed closely related haplotypes globally distributed [20], suggesting that they result from recent migrations associated with modern agriculture. Our sequencing of eight isolates of *M. incognita* and five of *M. javanica* from multiple continents has also shown that there is remarkably little genetic variation between isolates, and that this is true of both nuclear or mitochondrial genomes (Figure 3A; Supplementary Material, Figure S5A). The lack of intraspecific sequence diversity, even between samples taken from different continents, strongly suggests that agricultural environments do not contain indigenous populations but rather isolates that have only expanded with modern agriculture in the last few hundred years.

Nuclear sequence diversity exceeded mitochondrial diversity in comparisons between the MIG species (Figure 4). Although this is not typical of animals, the mitochondrial mutation rate is known to vary greatly among lineages [51,52]. It is, however, difficult to conclude that a low mutation rate alone is the cause of this extreme interspecific mitochondrial sequence identity since comparisons between the MIG and outgroups show much more variation in mtDNA than nuclear genomes (Figure 4). Between these closely related species the extremely high AT-content in the mitochondrial genome (84% for all MIG) may play a role. With mostly just two nucleotide character states out of four represented in the sequence, and most segregating intraspecific polymorphism being third codon position synonymous changes, saturation and homoplasy could be prevalent, leading to underestimation of the true divergence.

More genetic diversity was present within *M. arenaria* and within *M. floridensis* than the other MIG species examined, and other studies have also indicated that *M. arenaria* contains considerable diversity [53–55]. Our sampling for these two taxa was limited, and in order to understand the structure and level of diversity in the MIG species extensive geographic sampling for population genomics will be needed. Many additional RKN species are likely to fall within the MIG phylogenetic cluster based on classical and molecular characterization [24,26]. As we have shown above, phylogenomics of the nuclear genome is able to separate closely related species, and even low coverage sequencing without genome assembly will, given appropriate analyses, be able to place species robustly within this phylogeny. In turn, phylogenetic understanding can be transformed into insights into global dispersal patterns, adaptive evolution and plant host specialisation.

### Conclusions

We have sequenced the genomes of a diverse set of apomicts from the *Meloidogyne incognita* group of species and shown that the divergent genome copies, most likely from a hybridization event, predates the formation of these species. This group therefore shares the same parental species and evolutionary history. The MIG genomes are hypotriploid and subject to the action of non-crossover recombination both in homogenising the divergent loci, but also in generating distinctiveness between the genomes of different species. Non-crossover recombination is especially visible between the MIG genomes because of the divergent genomic copies present, and broader sampling of genomes will speak to its phenotypic importance in this important system. Our study demonstrates the power of comparative genomics to untangle biologically complex species histories and the data will provide an opportunity to study the complex evolution of sex and genome structure.

## Acknowledgements

This work was funded by the NERC through research grant NE/J011355/1 awarded to DL and MB. AS is funded through the ICA grant 031606a and the Israel Ministry of Science. DRL was funded through a James Hutton Institute postgraduate fellowship. We thank colleagues in Edinburgh Genomics and The Liverpool Centre for Genome Research for expert sequencing support. We thank Philippe Castagnone-Sereno, Etienne Danchin, Bob Robbins, Tom Prior for additional *Meloidogyne* samples, discussions, and advice. Christoph Hahn, Georgios Koutsovoulos and Sujai Kumar gave valuable bioinformatics advice. The University of Hull VIPER High Performance Compute cluster supported our analyses. We thank Africa Gómez for informative discussions and close reading of the manuscript.

## Methods

### Reproducibility

In order to make our analyses as reproducible as possible we provide a collection of Jupyter notebooks containing analysis scripts, specified parameters, descriptions, and details of the data versions used. It should be possible to re-create all figures from these notebooks. Phylogenomic analyses made use of ReproPhylo an environment for reproducible phylogenomics [56]. Raw data is published in BioProject PRJNA340324 and genome assemblies, intermediate data transformations, and methods notebooks can be found in the manuscript’s git repository: https://github.com/HullUni-bioinformatics/MIG-Phylogenomics#mig-phylogenomicsde50fe4.

An independent and permanent freeze of this git repository is published at doi:10.5281/zenodo.399475.

### Samples and Sequencing

The isolates chosen for sequencing, with information about their origins, and other metadata are given in Table S1 and the sequencing data including library size and number of reads is given in Table S2 (Also https://github.com/HullUni-bioinformatics/MIG-Phylogenomics, DOI:10.5281/zenodo.399475).

### Assembly and annotation

Sequence read quality filtering, *de novo* assembly, and genome annotation were carried out as detailed in the github repository. We used the Platanus assembler [28] because it is optimised for highly heterozygous genomes, and performed best in our hands with our data.

In addition to the five *de novo* reference assemblies (Table 1), protein coding gene data was created *via* read mapping to whole gene reference sequences (including introns and exons). For each sample, the reference gene sequences were taken from its conspecific reference genome. Quality trimmed read pairs were mapped to the gene dataset using the BWA package [57]. The resulting alignment was converted to fasta entries by creating a VCF file with FreeBayes [58], including SNPs and monomorphic positions. This VCF file, containing all the sequence positions, was then formatted as fasta. The resulting gene assembly was annotated using the protein2genome model in exonerate [59], to delineate intron-exon boundaries. For each sample, protein query sequences were taken from the conspecific reference genome (Also section 2 in https://github.com/HullUni-bioinformatics/MIG-Phylogenomics, DOI:10.5281/zenodo.399475).

### Orthology definition

We used OrthoFinder [32] to define orthology among the proteins obtained from the reference genomes and from the mapping assemblies. The analysis included all the available data for *M. floridensis*, *M. incognita*, *M. javanica*, *M. arenaria*, and *M. enterolobii*. Although currently OrthoFinder incorporates a phylogenetic step, the version we have used (git commit 2bb1fe3), did not include this feature and we thus created a phylogenetic filtering step (see below). We tested inflation values of 1.1 - 20, and selected the inflation value 2, as the setting resulting in the most orthology groups (OGs) containing at least one copy from each sample. For each resulting OG we reconstructed a gene tree, using a nucleotide MAFFT l-ins-i alignment [60] and alignment trimming which allowed up to 30% missing data and at least 0.001 similarity score (TrimAl, [61]). Alignments that contained two (or more) sequences from a single species that had a small or no overlap (< 20 bp) in the alignment were discarded to avoid erroneous consideration of two exons of the same orthologue as separate orthologues. The trees were reconstructed with the default parameters in RAxML [62]. Each gene tree was rooted with with *M. enterolobii*, and the ingroup was traversed to collapse sister leaves belonging to the same sample into a single leaf, keeping the least derived sequence out of the two, using ete2 [63]. Then the ingroup was clustered into two groups based on patristic distances, using a hierarchical clustering approach. OGs with more than one representative per sample in each cluster were discarded. OG contents were edited based on the results of the steps described above, and new alignments and gene trees were constructed. (Also section 3 in https://github.com/HullUni-bioinformatics/MIG-Phylogenomics, DOI:10.5281/zenodo.399475)

### Phylogenomic reconstruction and tests for phylogenetic conflicts within the data

We tested for the existence of conflicting phylogenetic signal in the nuclear protein coding gene trees. Conflicting signal is not unusual among nuclear gene trees, but in this study we were additionally exploring possible conflict within trees that might exist between the subtrees of the first and second homoeologs. In a first approach, we calculated a weighted Robinson-Foulds distance matrix [64] among all the gene trees and constructed a metric MDS plot to examine the number of gene tree clusters that existed in the data with the treeCl package [65]. In a second approach, we created 100 sets of randomized gene trees, and built a coalescent tree for each randomized set. In the randomization process we randomly assigned homoeolog identity to each homoeolog subtree, in each gene tree, such that for a given gene tree, subtree 1 would be denoted “homoeolog 1” in some of the randomized sets, and “homoeolog 2” in the other randomized sets, whereas subtree 2 would be called “homoeolog 2” and “homoeolog 1” respectively (Figure S4). We did not use the terms ‘genome A’ and ‘genome B’ when naming the homoeologs to indicate that we do not have synteny information for homoeologs from different orthology clusters. Using the suffixes ‘A’ and ‘B’ would erroneously create the impression that their genome copy assignment is known. The third approach was based on a maximum likelihood (ML) phylogenetic reconstruction of 100 randomized supermatrices. For each gene in a given randomized supermatrix, the first and second homoeologs were randomly denoted as homoeolog A and B, or vice versa, thus affecting the concatenation process. For example, in one supermatrix, sequences of the first homoeolog from gene 1 were concatenated with sequences of the first homoeolog of gene 2, while in another supermatrix the concatenation was inverted (Figure S3). The second and third approaches allowed us to test whether the two homoeologs had a shared phylogenetic history or distinctly different phylogenetic histories. We reasoned that if the two homoeologs had a shared organismal history and had co-evolved since the MIG ancestor this would be reflected in shared phylogenetic histories, and neither of the randomization schemes would affect the resulting topology. (Also section 4 in https://github.com/HullUni-bioinformatics/MIG-Phylogenomics, DOI:10.5281/zenodo.399475)

### Mitochondrial genome assembly, annotation, and tree reconstruction

An iterative mapping and extension approach was followed to assemble mitochondrial genomes. We used MITObim [66] to iteratively carry out a mapping assembly with the Mira assembler [67]. Genes from publically available mitochondrial genomes were used as seed sequences. Genes from the mitochondrial genomes of *M. incognita* (NC_024097), *M. javanica* (NC_026556) and *M. arenaria* (NC_026554) were used as seeds for the assembly of their conspecifics with mismatch cutoffs of 1 to 4. *M. incognita* (NC_024097) was also used as a reference for *M. floridensis, M. enterolobii and M. haplanaria*, albeit with a more relaxed mismatch cutoff (6 and 15 respectively). Mitochondrial genes were annotated with exonerate [59], using the protein2genome model and protein sequences from the reference mitochondrial genomes as queries. Mitochondrial ribosomal RNAs and the mitochondrial putative control regions were annotated with the est2genome model and the nucleotide sequences from the reference mitochondrial genome as queries. Two phylogenetic trees were reconstructed with mitochondrial genes. The first was rooted with sequences from *M. enterolobii* and the second was an unrooted tree that only included sequence from MIG species. The analysis was kept reproducible with ReproPhylo [56]. Single gene datasets were aligned with the l-ins-i algorithm in MAFFT [60] and trimmed with TrimAl [61] to exclude alignment columns with more than 10% missing data. Sequences were then concatenated into a supermatrix and a tree with reconstructed with RAxML [62] using the GTR-GAMMA model and allowing a separate model of evolution for each gene (Also see sections 5, 6 and 7 in https://github.com/HullUni-bioinformatics/MIG-Phylogenomics, DOI:10.5281/zenodo.399475.

### Coverage ratio between homoeologous contigs

We used contig and scaffold coverage data to identify regions of the genome that had evidence of being present in two copies rather than one. This approach is analogous to approaches to identifying sex chromosomes in heterokaryotypic organisms. In the case of the *Meloidogyne*, we were attempting to identify homoeologous contigs where one homoeolog was present in two copies while the other was present in one copy. We paired homoeolog contigs that carried shared orthologous loci, and created a set of 400 contig pairs for each of the apomict MIG reference genomes, ensuring the sets were homologous among the genomes. For each apomict MIG sample, trimmed reads were mapped to contig pairs from its conspecific genomes. For each contig pair, the coverage values were aligned according to the contig sequence pair alignment. Finally, per position coverage ratios were calculated (designating the contig with the highest coverage as the numerator) and the median ratio was inferred for each contig pair. For each sample we fitted functions with one and two gaussian components and computed the residual error in each case, to determine which of the two functions fit the distribution of median coverage ratio better. (Also section 9 in https://github.com/HullUni-bioinformatics/MIG-Phylogenomics, DOI:10.5281/zenodo.399475)

### Recombination

To quantify recombination we took a sliding window BLAST approach to characterise changing similarity profiles between sequences (Supplementary Methods, Figure S8A). For each sample pair, we collected long scaffolds from the query genome (>10 kbp). We then recovered the first two hits on the target assembly, for each window in the query scaffold. Windows were 5000 bp with 2500 bp overlap. We required matches to be at least 2500 bp long, and have BLAST E-values < 0.01. Cases in which two adjacent windows had the same two hits, but in reciprocal order were considered “events”, as long as the difference between the two hits was at least 7 mismatches, and that they existed on different scaffolds. Events that were also recovered when exchanging the query and the target genomes were considered crossover events, otherwise they were considered non-crossover events. We note that this criterion is not valid for *M. floridensis*, which is a mostly haploid genome assembly. Non-crossover event rates were expressed as the fraction of scaffolds with an event out of the total scaffold count. The Pearson correlation coefficient was calculated between the non-crossover event rates and tree distances, for samples in which at least 3000 long contigs (>10 kbp) were recovered. This analysis is detailed in section 10 in https://github.com/HullUni-bioinformatics/MIG-Phylogenomics, DOI:10.5281/zenodo.399475.

## Supplementary Materials and Methods

*Supplementary results* are found in the Supplementary Material file accompanying this manuscript and can be found at https://doi.org/10.6084/m9.figshare.4989173.v1. *Supplementary methods* are provided as a git repository under DOI https://doi.org/10.5281/ZENODO.399475. The jupyter notebooks containing all the analyses can be readily viewed in the live version of the repository at https://github.com/HullUni-bioinformatics/MIG-Phylogenomics/blob/master/README.md.

